# Redundant enhancers in the *iab-5* domain cooperatively activate *Abd-B* in the A5 and A6 abdominal segments of Drosophila

**DOI:** 10.1101/2021.05.22.445252

**Authors:** Nikolay Postika, Paul Schedl, Pavel Georgiev, Olga Kyrchanova

## Abstract

The homeotic *Abdominal-B* (*Abd-B*) gene belongs to *Bithorax* complex and is regulated by four regulatory domains named *iab-5, iab-6, iab-7* and *iab-8*, each of which is thought to be responsible for directing the expression of *Abd-B* in one of the abdominal segments from A5 to A8. It is assumed that male specific features of the adult cuticle in A5 is solely dependent on regulatory elements located in *iab-5*, while the regulatory elements in the *iab-6* are both necessary and sufficient for the proper differentiation of the A6 cuticle. Unexpectedly, we found that this long held assumption is not correct. Instead, redundant tissue-specific enhancers located in the *iab-5* domain are required for the proper activation of *Abd-B* not only in A5 but also in A6. Our study of deletions shows that the *iab-5* initiator is essential for the functioning of the *iab-5* enhancers in A5, as well as for the correct differentiation of A6. This requirement is circumvented by deletions that remove the initiator and most of the *iab-5* regulatory domain sequences. While the remaining *iab-5* enhancers are inactive in A5, they are activated in A6 and contribute to the differentiation of this segment. In this case, *Abd-B* stimulation by the *iab-5* enhancers in A6 depends on the initiators in the *iab-4* and *iab-6* domains.

**Summary Statement:** In *Drosophila*, the segmental-specific expression of the homeotic gene *Abdominal-B* in the abdominal segments is regulated by autonomous regulatory domains. We demonstrated cooperation between these domains in activation of *Abdominal-B*.

## Introduction

In *Drosophila melanogaster* segment identity in the posterior 2/3rds of body is controlled by the three homeotic genes, *Ultrabithorax* (*Ubx*), *abdominal-A* (*abd-A*) and *Abdominal-B* (*Abd-B*), which form the *bithorax* complex (BX-C) (Lewis, 1978). The specification of parasegments (PS)/segment identity depend upon the expression patterns of these three homeotic genes (Duncan, 1987; Karch et al., 1985; Karch et al., 1990; Maeda and Karch, 2015; Peifer et al., 1987). The genes are controlled by an array of nine regulatory domains, each of which is thought to be responsible for directing the expression of one of the homeotic genes in a spatio-temporal pattern appropriate for the particular PS/segment. The *Abd-B* gene is responsible for the specification and differentiation of PS10/A5, PS11/A6, PS12/A7, PS13/A8, in which the pattern of its expression is determined by four regulatory domains, *iab-5, iab-6, iab-7* and *iab-8* respectively (Fig. 1A).

**Fig. 1.**
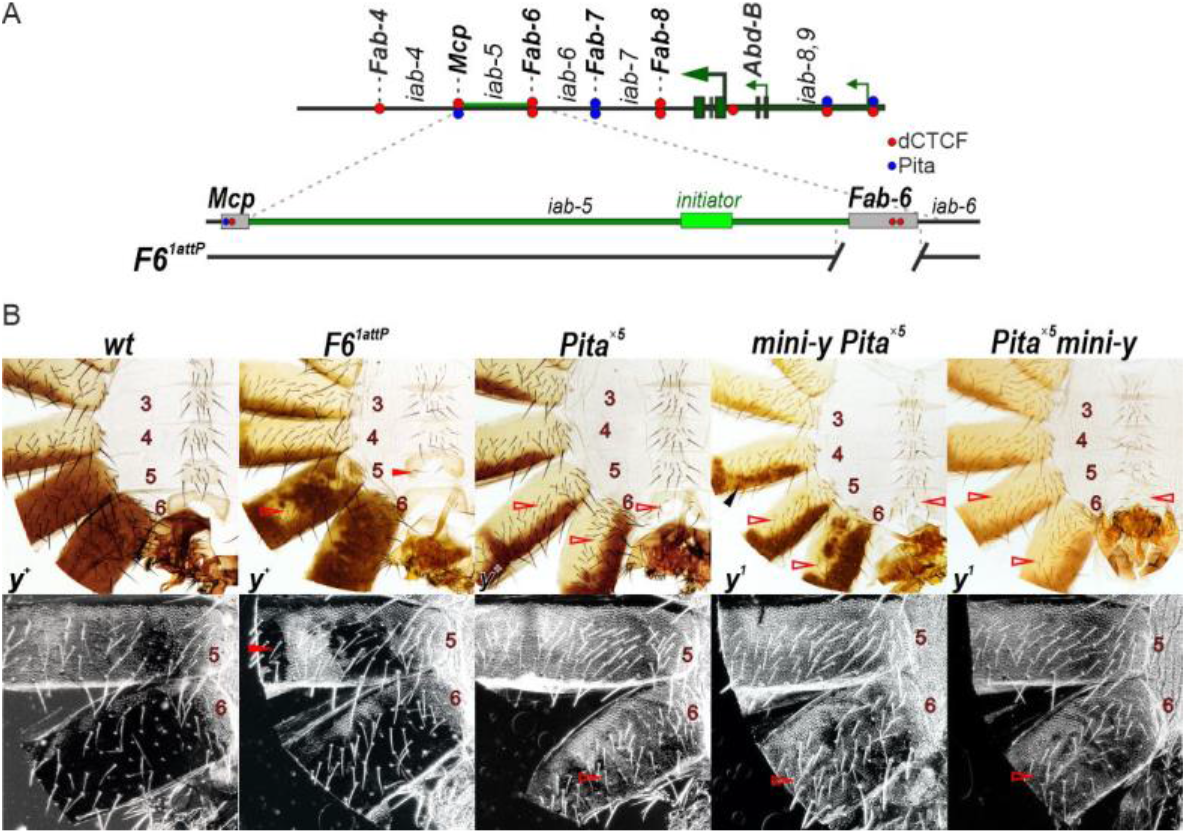
The substitution of the *Fab-6* boundary by Pita sites blocks the *Abd-B* expression in the A5 and A6 segments. (A) Scheme of the *Abd-B* regulatory region and the *F6*^*1attP*^ deletion. The *Abd-B* promoters are shown by green arrows. The dashed lines with colored circles mark boundaries. Pita and dCTCF are indicated by blue and red circles, respectively. The DNAse I hypersensitive sites of *Mcp* and *Fab-6* boundaries are shown as grey boxes. The endpoints of the *F6*^*1attP*^ deletion used in the replacement experiments are indicated by breaks in the black lines. (B) Morphology of the male abdominal segments (numbered) in *wt, F6*^*attP*^ and *Pita*^*×5*^ lines. In *Pita*^*×5*^ replacements males the A6 sternite has an intermediate form between quadrilateral (as in *wt* A5) and banana-like (as in *wt* A6) and is partially covered by bristles, while the tergite loses pigmentation and is covered by trichomes. The filled red arrowheads show morphological features indicative of GOF transformations. The empty red arrowheads show LOF transformations. Black arrowheads indicate pigmented spots that are induced by the *mini-y* expression. The localization of trichomes on the A5 and A6 tergites are shown in dark field.

Analysis of BX-C regulatory domains, including those controlling *Abd-B* indicate that they are composed of the same set of elements (Kyrchanova et al., 2015; Maeda and Karch, 2015). Each domain has an initiator element that sets the activity state (*on* or *off*) of the domain early in embryogenesis (Maeda and Karch, 2015; Mihaly et al., 2006; Peifer et al., 1987). Initiators responds to the maternal, gap and pair-rule gene products that subdivide blastoderm stage embryos along the antero-posterior axis into 14 parasegments (Busturia and Bienz, 1993; Casares and Sánchez-Herrero, 1995; Drewell et al., 2014; Ho et al., 2009; McCall et al., 1994; Qian et al., 1991; Shimell et al., 1994; Starr et al., 2011). For example, in PS10/A5, the *iab-5* initiator turns *on* the *iab-5* domain, while the adjacent *iab-6* and other more distal (relative to centromere) domains remain in the *off* state (Iampietro et al., 2010). In PS11/A6, the initiator in *iab-6* turns the domain *on*. While *iab-5* is also active in PS11, *iab-7* and *iab-8* are *off*. The gene products responsible for setting the activity state of the BX-C domains disappear during gastrulation and different mechanisms are deployed to remember the *on* or *off* state. The *on* state is maintained by Trithorax group proteins, while the *off* state is maintained by Polycomb group proteins (Busturia and Bienz, 1993; Kassis et al., 2017; Kuroda et al., 2020; Simon et al., 1992; Shimell et al., 2000; Ciabrelli et al., 2017; Müller and Bienz, 1992). These factors interact with special elements in each domain called Trithorax or Polycomb response elements (TREs or PREs). Finally, each domain has a stage and tissue specific enhancers which are responsible for activating patterns of homeotic gene expression that drive PS/segment differentiation (Maeda and Karch, 2015). Each domain is bracketed by chromatin boundary elements (Barges et al., 2000; Bender and Lucas, 2013; Bowman et al., 2014; Galloni et al., 1993; Gyurkovics et al., 1990; Hagstrom et al., 1996; Karch et al., 1994; Kyrchanova et al., 2020; Mihaly et al., 2006, 200; Zhou et al., 1996). The boundaries in the *Abd-B* region (*Fab-6, Fab-7* and *Fab-8*) have two important functions. The first is to block crosstalk between adjacent regulatory domains so that they can function autonomously. The loss of one of these boundaries leads to the ectopic activation/silencing of neighboring regulatory domains. For example, deletion of the *Fab-6* boundary element can result in the ectopic activation of *iab-6* and silencing *iab-5* in PS10/A5 leading respectively to gain-of-function (GOF) and loss-of-function (LOF) transformation of PS/segment (Iampietro et al., 2010; Postika et al., 2021). The second function is boundary bypass that enables enhancers in the *Abd-B* regulatory domains to bypass intervening boundaries and activate *Abd-B* (Kyrchanova et al., 2019a; Kyrchanova et al., 2019b; Postika et al., 2018).

While identity of PS10-PS13/A5-A8 is determined by the pattern of *Abd-B* expression in both sexes, the phenotype of the adult cuticle in segments A5 and A6 in *Drosophila melanogaster* differs in males and females (Jeong et al., 2006; Kopp et al., 2000; Massey and Wittkopp, 2016; Williams et al., 2008). In females, cuticle pigmentation and morphology in A5 and A6 are similar to that in more anterior segments whose identity is determined by *abd-A*. In these segments the tergite has a posterior stripe of dark pigmentation, while the sternite has a quadrilateral shape and has multiple bristles. The pigmented stripe in tergites A2-6 is generated by the *yellow* (*y*) and *tan* genes, which are regulated by the *optomoter blind* (*omb*) gene (Kopp and Duncan, 1997). The *bric-a-brac* (*bab*) complex encodes DNA-binding proteins that repress the expression of the genes responsible for cuticle pigmentation (Couderc et al., 2002; Kopp et al., 2000; Roeske et al., 2018). While female pupae express *bab* in abdominal segments A2–A6, *bab* expression in males is limited to segments A2–A4. The sex specific pigmentation pattern and cuticle morphology in A5 and A6 in males depend upon *Abd-B* and the male product of the *double-sex* gene (*dsx*^*M*^), which together function to repress expression of *bab* genes in cells giving rise to the A5 and A6 (Kopp et al., 2000; Massey and Wittkopp, 2016; Wang et al., 2011).

The level of *Abd-B* expression in PS10/A5 and PS11/A6 is not the same and correlates with their distinctive morphology. The *Abd-B* expression in A5 is relatively low and this segment has morphological features of the A4 where *abd-A* is expressed: the A5 sternite has a quadrilateral shape and has multiple bristles, while the A5 tergite is covered by small trichome hairs. However, due to the expression of *Abd-B* in the A5 segment of males, differences are observed: sternite becomes wider, tergite is completely pigmented, and trichomes are less dense (Celniker et al., 1990; Maeda and Karch, 2015) (Fig.1). The higher levels of *Abd-B* in A6 are accompanied by specific morphological features in both the sternite and tergite. The A6 sternite lacks bristles and has a unique ‘banana’ shape, while the trichomes on the fully pigmented tergite are restricted to the anterior and dorsal margins instead of covering nearly the entire tergite.

Here, we have investigated the mechanisms responsible for regulating *Abd-B* expression during the differentiation of the male cuticle in segments A5 and A6. We have found that *iab-5* and *iab-6* domains share a common set of partially redundant cuticle enhancers located in *iab-5* that are critical for male specific differentiation of the cuticle of A5 and A6 segments.

## Results

### Inactivation of the *iab-5* domain affects expression of *Abd-B* in the A6 segment

In contrast to the boundaries of the *Abd-B* region, that have blocking and bypass activity (Kyrchanova et al., 2016; Kyrchanova et al., 2019a; Postika et al., 2018) the heterologous boundaries, for example, DNA fragment consisting of five binding sites for the multi-Cys2His2 zinc finger protein Pita (*Pita*^*×5*^), have only blocking activity. To test whether the regulation of *Abd-B* by *iab-5* requires the bypass boundary activity, we took advantage of *F6* ^*1attP*^ replacement platform, in which a 1389 bp sequence spanning the *Fab-6* boundary was substituted by an *attP* site (Postika et al., 2021). We have replaced *Fab-6* with *Pita*^*×5*^. In order to assess the activity of cuticle enhancers in *iab-5* and *iab-6*, we included a *mini-yellow* (*mini-y*) reporter which we placed either upstream of *Pita*^*×5*^ or downstream so that it would be located in the *iab-5* (*mini-y Pita*^*×5*^) or *iab-6* (*Pita*^*×5*^ *mini-y*) domains, respectively (Fig. S1). The reporter consists of a *yellow* (*y*) cDNA fused to the 340 bp *y* promoter. As it lacks the enhancers of the endogenous *y* gene, its expression depends upon nearby enhancers. Expression of *mini-y* was examined in a *y*^*1*^ background. In flies carrying the null *y*^*1*^ allele, the *tan* gene is appropriately expressed in A5 and A6 reflecting the *Abd-B* activity, and the resulting pigmentation in the tergite is light brownish, not black (Camino et al., 2015; Rebeiz and Williams, 2017).

*F6*^*1attP*^ deletion mutant males differ from *wild type* (*wt*) in that segment A5 has an incomplete GOF and LOF transformation (Fig. 1). The A5 sternite has a shape like that normally observed in A6, but with several bristles, the A5 tergite has patches of cuticle that lack trichomes, which is indicative of a GOF transformation towards A6 identity. On the other hand, large portions of the A5 cuticle also lack *tan* pigmentation, indicating that the cells have an A4 identity. There are also LOF phenotypes in A6 including bristles on the sternite and regions of the tergite that are depigmented or have ectopic trichomes.

In the *Pita*^*×5*^ replacements with or without *mini-y* the GOF transformations of A5 are eliminated (Fig. 1). Since *Pita*^*×5*^ does not support bypass, A5 resembles A4: the sternite has a quadrilateral shape, and is covered in bristles, while the tergite is covered in trichomes, and instead of being covered in pigmentation, there is only a posterior stripe. This result shows that *iab-5* is unable to activated *Abd-B* in A5, and that the cells in this segment have an A4 identity. However, there is also an unexpected result: the differentiation of A6 is altered compared to *wt*. The defects are most clearly evident when *mini-y* is excised and *y*^*+*^ allele is introduced. Fig. 1 shows that pigmentation of the A6 tergite resembles A4: there is only a stripe of pigment along the posterior margin of the tergite. In addition, the A6 tergite is covered in trichomes just like A4. While the A6 sternite has a nearly normal shape, there are multiple bristles. These LOF transformations indicate that the *Pita*^*×5*^ insulator disrupts *Abd-B* dependent cuticle differentiation not only in A5, but also in A6. These results suggest that *iab-5* is required for the proper activation of *Abd-B* in the cuticle in both A5 and A6.

Further support for this conclusion comes from analysis of *mini-y* expression in the *y*^*1*^ background. In *Pita*^*×5*^*mini-y* males the reporter located in the *iab-6* domain it is not turned on in neither in A5, nor in A6. Instead, only the *tan* gene is expressed in an A4 like pattern. This result would indicate that enhancers in *iab-5* are required to drive expression of *mini-y* inserted in *iab-6* in the A6 tergite. When the *mini-y* reporter is in *iab-5* we observe a mosaic pattern of pigmentation in the posterior stripes of the A4, A5 and A6 segments.

### The *iab-5* domain contains a set of redundant cuticle enhancers that can drive *yellow* expression outside BX-C

To map enhancers in the *iab-5* regulatory domain responsible for *Abd-B* expression, we linked 1-3 kb overlapping DNA sequences from the *iab-5* domain (*i5*^*1*^ *– i5*^*7*^ and *i5*^*ini*^) to a *y* reporter in a transgene that also carries a *mini-white* (*w*) (Fig. S2). To reduce potential position effects, we placed *Pita*^*×5*^ upstream of the *iab-5* DNA fragments. Using *phiC31*-mediated recombination (49), we integrated a collection of eight *i5* transgenes into a well characterized 86Fb platform. Of these *i5* fragments, only three, *i5*^*1*^ (1013 bp), *i5*^*2*^ (2145 bp) and *i5*^*7*^ (2524 bp), activated *y* expression in the cuticle. For all three, pigmentation was observed in the A5 and A6 tergites. Interestingly, we found that the *i5*^*7*^ fragment was only able to activate *y* in the forward (genomic) orientation. We tested two *i5*^*7*^ sub-fragments from the proximal (*i5*^*S5*^) and distal (*i5*^*S6*^) ends relative to the centromere. Of these, only *i5*^*S6*^, activated *y*. Thus, in larger *i5*^*7*^ fragment, the enhancer in *i5* ^*S6*^ must be located next to the promoter to function. Since the *i5*^*1*^ and *i5*^*2*^ overlapped, it seemed possible that they share the same enhancer. To test this, we generated three smaller fragments (*i5*^*S1*^, *i5*^*S2*^, and *i5*^*S3*^) spanning most of *i5*^*1*^ and *i5*^*2*^. Of these only *i5*^*S2*^ which includes the overlap between *i5*^*1*^ and *i5*^*2*^ activates *mini-y*.

### Functioning of the *iab-5* enhancers in the *iab-6* domain

We next determined whether *iab-5* sequences are able to regulate *Abd-B* in A6 when placed in *iab-6*. We used the *F6*^*1attP*^ landing platform to insert the same collection of *iab-5* sequences into the *iab-6* regulatory domain (Fig. S3). The starting transgene included *Pita*^*×5*^ to block crosstalk with *iab-5* and excisable *mCherry* and *mini-y* reporters arranged so that they are located in *iab-6* in the replacements (Fig. 2, Fig. S1).

**Fig. 2.**
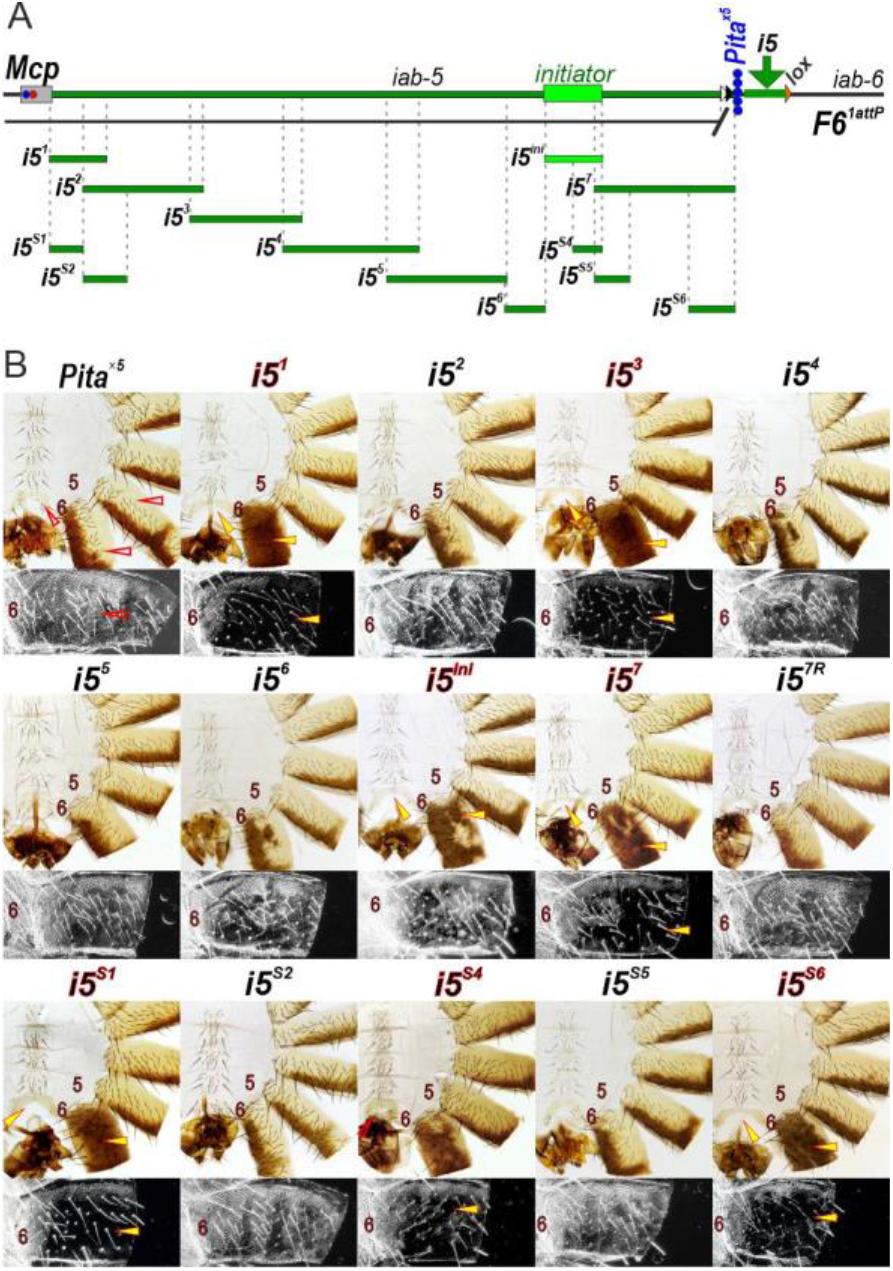
Testing regions in the *iab-5* domain that stimulate *Abd-B* expression in the A6 segment. (A) Scheme of *iab-5* with the Pita^×5^ replacements in the *F6*^*1attP*^ platform. The *i5* fragments tested for enhancer activity are shown as green lines, the *i5*^*ini*^ fragment including the initiator is shown as light green line. The test fragments were integrated near *Pita*^*×5*^ (five blue circles vertically) in the *iab-6* domain. (B) Morphology of male abdominal segments in transgenic lines with different *Pita*^*×5*^*-i5* substitutions. The localization of trichomes in the A6 tergite is shown in dark field. The yellow arrowheads show the signs of rescue the LOF phenotype in A6. All other designations are the same as in Fig. 1.

Three of the *iab-5* sequences, *i5*^*1*^, *i5*^*3*^, and *i5*^*7*^ are able to stimulate *mini-y* expression to different extents in the A6 segment (Fig. S3). While both *i5*^*1*^ and *i5*^*7*^ also stimulated *y* when inserted in 86Fb platform, *i5*^*3*^ did not. Conversely, *i5*^*2*^ failed to function when placed in *iab-6*, while it is active in the 86Fb platform. It seems likely that “position effects” are responsible for the differences in the activity of the *iab-5* sequences when linked to the *y* reporter in 86Fb or inserted in *iab-6*. As would be expected from their placement in *iab-6, i5*^*1*^, *i5*^*3*^ and *i5*^*7*^ do not activate the reporter in more anterior segments. Surprisingly, insertion of the *i5*^*7*^ fragment in the reverse orientation (*i5*^*7R*^) stimulates *mini-y* expression in posterior stripes not only in the A6 tergite, but also in the A5 and A4 tergites (Fig. S3). Since the distal part of *i5*^*7*^, *i5*^*S6*^, induces a much stronger activation of the *mini-y*, it would appear that sequences elsewhere in *i5*^*7*^ contain a silencer.

As the reporters compete with *Abd-B* for enhancer activity we assessed the cuticle phenotypes after removing the reporters and introducing a *y*^*+*^ allele (Fig. 2). When *i5*^*1*^, *i5*^*3*^ or *i5*^*7*^ are included in the *Pita*^*×5*^ replacements the phenotype of A6 is close to *wt*. Even though *i5*^*S2*^ and *i5*^*2*^ activate *mini-y* at 86Fb, neither could rescue the LOF phenotypes induced by *Pita*^*×5*^. On the other hand, *i5*^*S1*^ and *i5*^*3*^, which does not stimulate *mini-y* at 86Fb, completely rescues the *Pita*^*×5*^ induced LOF phenotypes in A6 (Fig. 2). The remaining fragments that are active when introduced into *iab-6* are *i5*^*ini*^ and *i5*^*7*^. The former is not active at 86Fb, while the latter is. Both partially rescue the *Pita*^*×5*^ induced defects in A6. The fragment of *i5*^*ini*^, *i5*^*S4*^, restore A6 identity with variable efficiency. As was the case in 86Fb, the *i5*^*7*^ enhancer activity is orientation dependent and it is not observed in *i5*^*7R*^. Thus, there are several enhancers in *iab-5* that could help drive *Abd-B* expression in the cuticle and generate morphological features that are characteristic of A6.

These results suggest that enhancers in *iab-5* are important for the proper differentiation of the adult cuticle in A6.

### Deletion of the *iab-5* initiator disrupts morphology of the A5 and A6 segments

According to the previous results deletion of the *iab-5* initiator will disrupt the development of the adult cuticle not only in A5 but also in A6. We used CRISPR/Cas9 to delete a 1975 bp genomic DNA segment that spans the *iab-5* initiator and replace it with an *attP* site and an excisable *dsRed* reporter under control of the *3×P3 hsp70* promoter (Fig. S1). As expected for an initiator deletion, the A5 segment in *i5*^*attP*^ males resembles A4 (Fig. 3, Fig. S4 and S5). Critically, this is not the only phenotypic alteration in *i5*^*1attP*^ males: the A6 sternite has an intermediate quadrilateral shape and also has bristles, while the A6 tergite has an irregular and variable pigmentation. In addition, trichome hairs are found in large patches often coinciding with areas of depigmentation. These results show that deletion of the *iab-5* initiator affects *Abd-B* expression in both the A5 and A6 segments. To confirm that *iab-5* is not properly activated in *i5*^*1attP*^ we integrated a *mini-y* reporter using the *attP* site. As expected, the *mini-y* reporter introduced into *iab-5* is *off* in A5. In A6, black pigmentation is restricted to several patches on the tergite, while *tan*-only dependent pigmentation occupies a somewhat larger area (Fig. 3, Fig. S5).

**Fig. 3.**
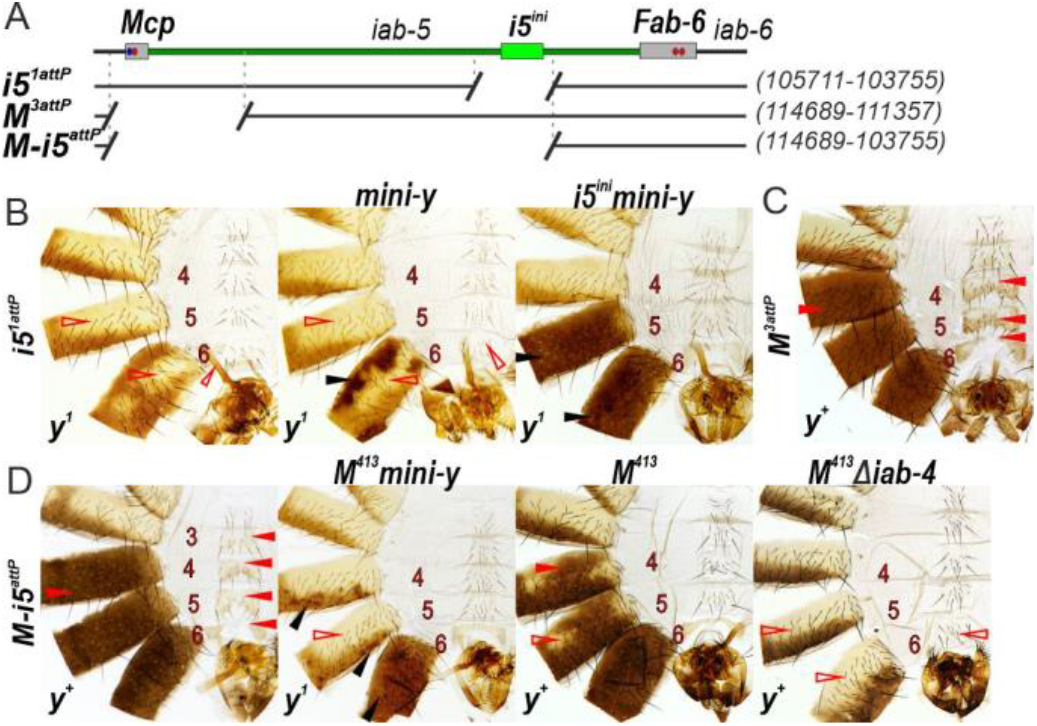
Deletions in the *iab-5* and *iab-4* domains. **(A)** Scheme of the *i5*^*1attP*^, *M*^*3attP*^ and *M-i5*^*attP*^ deletions. The endpoints of the deletions are indicated by breaks in the black lines. The coordinates of endpoints are according to the complete sequence of BX-C in SEQ89E numbering (Martin et al., 1995). Morphology of the male abdominal segments in transgenic line carrying (B) the *i5*^*1attP*^ deletion with (*y*^*1*^; *i5*^*1attP*^*mini-y*) or without (y^1^; *i5*^*1attP*^) the *mini-y* reporter or with re-integration of the 1019 bp *iab-5* initiator and the *mini-y* reporter (*i5*^*ini*^*mini-y*); (C) the *M*^*3attP*^ platform; (D) the *M-i5*^*attP*^ platform, integration of the *M*^*413*^ insulator in *M-i5*^*attP*^ with (*M*^*413*^*mini-y*) or without (*M*^*413*^) *mini-y*, deletion of the *iab-4* region in *M*^*413*^ (*M*^*413*^*Δiab-4*). Designations are the same as in Fig. 1.

To confirm that the observed effects on *mini-y* and A6 morphology are induced by deletion of the *iab-5* initiator, we introduced a 1025 bp *i5*^*ini*^ fragment together with *mini-y* into *i5*^*1attP*^. The resulting flies have *wt* morphology except for 1-2 bristles on the A6 sternite, and the *mini-y* reporter is expressed throughout the tergite in A5 and also A6 (Fig. 3). The presence of bristles on the A6 sternite is likely due to competition between the *mini-y* and *Abd-B* promoters.

### Deletions of the *iab-5* regulatory domain: impact on segment specification

To further assess the functional role of the *i5* enhancers in both A5 and A6, we deleted most of *iab-5* sequence and then reintroduced *i5* enhancers in various combinations. For this purpose, we used Cre-mediated recombination between *lox* sites located in the *i5*^*attP*^ and an *Mcp* boundary deletion, *M*^*3attP*^, in which a 3,333 bp sequence spanning the region around the *Mcp* boundary was substituted by *attP* and *lox* sites (Fig. 3, Fig. S6 and S7). After Cre recombination, the final deletion, *M-i5*^*attP*^, is 10935 bp. It extends from the centromere proximal side of the *Mcp* boundary through the *iab-5* initiator, leaving the 2126 bp *i5*^*7*^ sequence (Fig.2) and single *attP* and *lox* sites. *M-i5*^*attP*^ males have a pigmented A4 segment and display other signs of GOF transformation of not only A4 and A5, but also A3: the sternites have two lobes somewhat like the A6 sternite, while there is a depletion of the trichomes on the tergites (Fig. 3, Fig. S4).

Aiming to prevent the *iab-4* domain from activating *Abd-B*, we re-introduced a minimal *M*^*413*^ boundary, characterized previously (Kyrchanova et al., 2007), with the *mini-y* reporter using the *phiC31* integration system (Fig. S1). The resulting *M*^*413*^*mini-y* replacement contains only the *i5*^*7*^ sequence. As would be expected since there is no initiator in *iab-5*, the domain is inactive in A5 and *mini-y* is not expressed in this segment. However, in spite of the fact that the *iab-5* domain is inactive, the phenotype of A6 resembles *wt* and the *mini-y* reporter, which is located in the inactive *iab-5* domain, is expressed throughout the A6 tergite (Fig. 4). When the reporters are excised, the minimal *Mcp*^*413*^ boundary is not able to prevent *iab-4* from activating the enhancer in *i5*^*7*^, or *Abd-B* directly. In addition to having a *wt* A6 segment, the tergites in A4 and A5 are nearly covered in pigmentation indicating that the *Abd-B* gene active in both of these segments (Fig. S8).

**Fig. 4.**
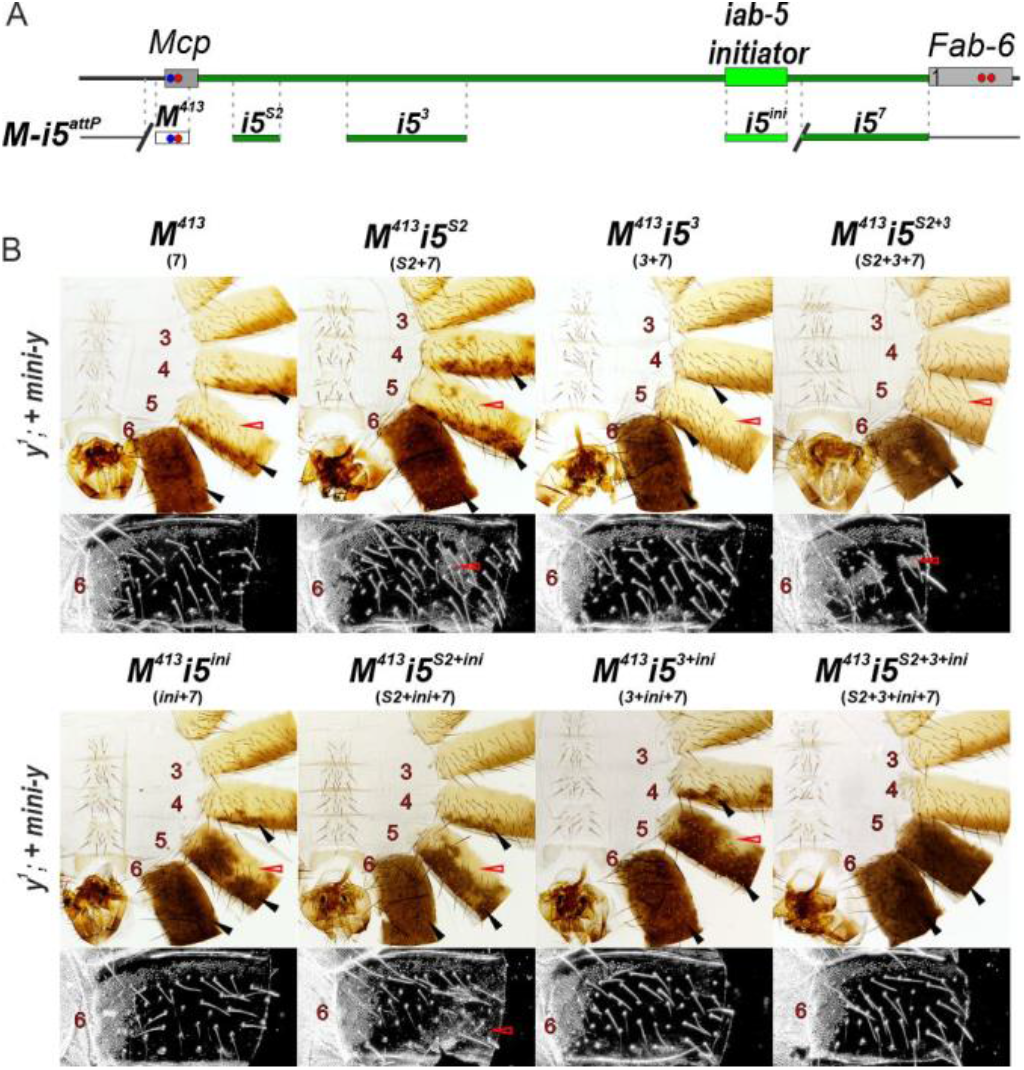
Reconstruction of the *iab-5* domain with *i5* fragments integrated in the *M-i5*^*attP*^ platform. (A) Scheme of the *M-i5*^*attP*^ platform and derivative lines carrying insertion of different *i5* combinations with the *M*^*413*^. (B) Morphology of the male abdominal segments in transgenic line carrying the *M-i5*^*attP*^ and different combination of *i5* fragments with the *M*^*413*^. In all transgenic lines the *mini-y* and *mCherry* reporters are present. Designations are the same as described in Fig. 1.

To test the role of the *iab-4* regulatory region in the GOF transformation of A4 in *Mcp*^*413*^, we deleted a 4,401 bp sequence (*iab-4Δ*) including the *iab-4* initiator as described previously (Postika et al., 2018). The deletion of these *iab-4* sequences not only reverts the GOF transformations in *M*^*413*^, but also results in a dramatic LOF transformation of *both* A5 and A6 (Fig. 3). While A5 resembles A4 in *M*^*413*^*iab-4Δ* males, the pigmentation patterns in the A6 tergite range from a few dark spots to almost ubiquitous pigmentation (Fig. S9). The A6 sternite is also misshapen and covered in bristles. Since A6 appears *wt* when the *iab-4* domain is intact, it would appear that in the *M*^*413*^ platform sequences in *iab-4* are able to collaborate with elements in *iab-6* to activate enhancers in *i5*^*7*^, and direct the proper expression of *Abd-B* in A6.

### Reconstructing a minimal *iab-5* domain

We next used the *M-i5*^*attP*^ platform to reconstruct a minimal *iab-5* regulatory domain. Since *Mcp*^*413*^ in combination with the two reporters is more effective in insulating against elements in *iab-4*, we will first consider the functioning of different *iab-5* sequences in the presence of the reporters. In the first set of experiments, we tested *i5*^*S2*^ and *i5*^*3*^. Since the *M-i5*^*attP*^ deletion retains the *i5*^*7*^, it is included in all of the replacements we tested. Thus, the three combinations are *i5*^*S2*^+*i5*^*7*^; *i5*^*3*^+*i5*^*7*^ and *i5*^*S2*^+*i5*^*3*^+*i5*^*7*^ (Fig. 4). In both *i5*^*7*^ and *i5*^*S2*^+*i5*^*7*^ the *mini-y* reporter is expressed in a mosaic pattern along the posterior margin of A4 and A5. In contrast, we observed only rare spots of dark pigmentation in the A5 segment in combinations containing *i5*^*3*^. Thus, the *i5*^*3*^ region has a negative effect on *mini-y* expression in *cis*. In all three combinations, the anterior 2/3rds of A5 tergite is largely devoid of pigmentation, indicating that the *tan* gene is also not expressed in much of the tergite. At the same time, the A6 segment has a nearly *wt* phenotype.

We next tested the same combinations of *i5* enhancers with the initiator, *i5*^*ini*^. The *i5*^*ini*^+*i5*^*7*^ combination expands the expression domain of *mini-y* in A5, while having minimal effect on expression in A4. However, there are regions in the anterior of the A5 tergite where *mini-y* is not expressed (Fig. 4). While adding *i5*^*S2*^ has little effect on the pattern of *mini-y* expression (*i5*^*S2*^+*i5*^*ini*^+*i5*^*7*^), there is a noticeable expansion in the expression area in A5 when *i5*^*3*^ is combined with the initiator (*i5*^*3*^+*i5*^*ini*^+*i5*^*7*^) (Fig. 4). This is the opposite of what was observed for *i5*^*S2*^+*i5*^*S7*^ and *i5*^*3*^+*i5*^*S7*^combinations without the initiator sequence. However, even in this case, *mini-y* expression is not observed throughout the A5 tergite. On the other hand, when the initiator is combined with all three sequences (*i5*^*S2*^+*i5*^*3*^ + *i5*^*ini*^+*i5*^*7*^), *mini-y* is expressed throughout the entire A5 tergite as is *tan*, while the ectopic activation in A4 is absent. (Fig. 4). Thus, this combination appears to be sufficient for full domain function.

We also examined the activity of the *iab-5* enhancers after reporter excision (Fig. S8). The GOF transformations (misshapen sternites and loss of trichome hairs) in the morphology of segments A3-6 in the starting *M-i5*^*attP*^ platform are largely rescued by the introduction of the *Mcp* boundary, *M*^*413*^. However, as mentioned above, the pigmentation patterns in A4 and A5 are still abnormal. The former has patches of ectopic pigmentation, while the latter is not fully pigmented. The pigmentation patches in anterior of A4 and A5 mostly disappear in the *i5*^*S2*^+*i5*^*7*^ combination. When *i5*^*S2*^+*i5*^*7*^ are combined with *i5*^*3*^ there is a further suppression in A5 pigmentation, and a loss of pigmentation in A4. Thus, the *i5*^*3*^ and *i5*^*7*^ sequences, in cooperation *i5*^*S2*^, can block the activation of *Abd-B* expression mediated by sequences in the *iab-4* domain. However, as was observed when *mini-y* is present, the *iab-5* enhancers in *i5*^*S2*^, *i5*^*3*^ and *i5*^*7*^ are unable to direct the proper development of A5 unless the *iab-5* initiator is also present. Addition of *i5*^*ini*^ to *i5*^*7*^, or *i5*^*S2*^+*i5*^*7*^, or *i5*^*3*^+*i5*^*7*^ substantially expands the area of pigmentation not only in the A5 tergite, but also in A4 (Fig. S6). As was observed for the *mini-y* reporter, combining *i5*^*ini*^ with *i5*^*S2*^+*i5*^*S3*^+*i5*^*S7*^ gives what appears to be a fully *wt* pattern of pigmentation in both A4 and A5. Thus, the *i5*^*S2*^+*i5*^*S3*^+*i5*^*S7*^ combination blocks incorrect activity of the *iab-4* and *iab-5* initiators in the A4 segment and is sufficient in cooperation with *i5*^*ini*^ for the proper stimulation of *Abd-B* in the A5 segment.

## Discussion

In *Drosophila melanogaster*, segments A5 and A6 in males are completely pigmented, while the cuticle morphology of A6 is distinctive from that in more anterior segments. Unexpectedly, we find that the regulatory elements in *iab-6* are not in themselves sufficient to direct the proper differentiation of the cuticle in males. Instead, the *iab-5* domain contains several enhancers that are required for male pigmentation not only in A5 but also A6. These *iab-5* enhancers also play a role in generating the proper distribution of trichome hairs in the A6 tergite, and in ensuring that the sternite is devoid of bristles. According to the generally accepted model (Maeda and Karch, 2015), the most likely initiator function is turning the *iab-5* domain *on* so that the cuticle enhancers are active. Deletion of the *iab-5* initiator results in a transformation of A5 into A4. We find that the *iab-5* initiator is also required for the proper differentiation of A6.

Functional dissection of *iab-5* suggests that at least four distinct DNA sequences (*i5*^*S2*^, *i5*^*3*^, *i5*^*7*^ and the *iab-5* initiator *i5*^*ini*^) are required for proper differentiation of A5. With respect to the cells in the cuticle in which these enhancers are active, it would appear that they have overlapping rather than completely distinct activities. Several lines of evidences (orientation or position dependent activation of the *yellow* reporter) suggest that the *i5*^*S2*^, *i5*^*3*^ and *i5*^*7*^ sequences contain not only tissue-specific enhancers but also silencers. The *i5* sequences also seem to help the *Mcp*^*413*^ boundary block interactions between the *iab-4* and *iab-5* initiators/regulatory elements.

While our results demonstrate that the proper differentiation of A6 depends upon enhancers in *iab-5* to drive *Abd-B* expression in the appropriate manner, previous studies showed that in addition to the initiator the *iab-6* domain has at least two regions that are required for the proper differentiation the A6 segment in the males (Iampietro et al., 2010). The deletion of the *iab-6* initiator results in a clean LOF transformation of A6 into A5. Thus, the *iab-5* enhancers that appear to complement the cuticle enhancers in *iab-6* are not able to function in A6 unless the enhancers in *iab-6* are also active and can communicate with *iab-5*. Further studies will be needed to refine and extend these findings.

## Methods

### Generation of *i5*^*1attP*^, *M*^*3attP*^ and *M-i5*^*attP*^ platforms

The deletions were obtained by CRISPR/Cas9 method (Fig. S1). As a reporter, we used *pHD-DsRed* vector (Addgene plasmid # 51434). The plasmid was constructed in the following order: *proximal arm-attP-lox-3×P3:DsRed-SV40polyA-lox-distal arm*. Arms were amplified by PCR from DNA isolated from *Oregon* line. For generation of the *i5*^*1attP*^ deletion, homology arms were obtained by DNA amplification between primers: TGTCGAGGTCCCGAAATG and ACGTCACTTGGCTGAAATGC; CAGACAGGTCCATCGGGG and TTGTTGAGGGTTGGTTGTG. For ^*M3attP*^: ATAACTAGTCCTAAATTACGACCACGAC and ATACTCGAGCCCATAAACAGCACGGC; ATAGCGGCCGCATTTTAATCGAGCCATC and CGAGAATTCCTAGAATGAGTAG. The guide RNAs were selected using the program “CRISPR optimal target finder” (O’Connor-Giles Lab). For *i5*^*1attP*^ deletion: TTTCGGGACCTCGACACGTT_TGG and TTGGCCCCGATGGACCTGTC_TGG. For *M*^*3attP*^: CACTGACAGAGTCAGGCTCG_TGG and CATACTTGCCCCGTACTTGC_CGG. The breakpoints of the designed deletion: *i5*^*1attP*^ -3R:16877730.. 16879686 (1957 bp) and *M*^*3attP*^ - 3R:16872084..16868751 (3333 bp), according Genome Release r6.36.

To generate the deletions, the plasmid construct was injected into embryos: *y*^*1*^ *M{Act5C-Cas9*.*P*.*RFP-}ZH-2A w*^*1118*^ *DNAlig4[169]* (BL 58492 stock, Bloomington Drosophila Stock Center) together with two gRNAs. The F0 progeny were crossed with *y w*; *TM6/MKRS* flies. Flies with potential deletions were selected on the basis of dsRed-signal in the posterior part of their abdomens and these flies were crossed with *y w*; *TM6/MKRS* flies. All independently obtained flies with *dsRed* reporter were tested by PCR. The successful deletions events were confirmed by sequencing of PCR products. Next, *dsRed* reporter was deleted by Cre/*lox* recombination.

To create *M-i5*^*attP*^ (Fig. S4) the *i5*^*1attP*^ and *M*^*3attP*^ were crossed with line expressing Cre recombinase (#1092, Bloomington Drosophila Stock Center). Then, *i5*^*1attP*^/+; *CyO, P{w[+mC]=Crew}DH1/+* was crossed with *M*^*3attP*^/+; *CyO, P{w[+mC]=Crew}DH1/+*. Next, the *i5*^*1attP*^/*M*^*3attP*^; *CyO, P{w[+mC]=Crew}DH1/+* males and females were crossed with each other and male offspring with the expected phenotypes were crossed with *y w*; *TM6/MKRS* flies. The deletion was confirmed by PCR and sequencing.

### Generation of transgenic lines carrying different insertions in the *attP*-platforms

The replacement vector was a plasmid with the *mini-yellow* and *mCherry* reporters as shown in Fig. S1. The *iab-5* fragments were obtained by PCR amplification. Their coordinates are: *i5*^*1*^: 112812-113529; *i5*^*2*^: 111101-113245; *i5*^*3*^: 109349-111346; *i5*^*4*^: 107265-109694; *i5*^*5*^: 105709-107851; *i5*^*5*^: 105030-105750; *i5*^*7*^: 101629-104152; *i5*^*ini*^: 104011-105035; *i5*^*S1*^: 113227-113824; *i5*^*S2*^: 112455-113245; *i5*^*S3*^: 111607 - 112829; *i5*^*S4*^: 104016 - 104537; *i5*^*S5*^: 103516 - 104152; *i5*^*S6*^: 101629 - 102685, according to the published sequences of the Bithorax complex (Martin et al., 1995).

Integration of the plasmids in the landing platforms was achieved by injecting the plasmid and a vector expressing the *фC31* recombinase into embryos of *yw; i5*^*1attP*^*/i5*^*1attP*^, or *yw; M*^*3attP*^/*M*^*3attP*^, or *yw; M-i5*^*attP*^*/M-i5*^*attP*^ lines. The successful integrations were selected on the basis of expression of *mini-y* in abdominal segments. The integration of the replacement DNA fragments was confirmed by PCR. The *yellow* and *mCherry* reporters were excised by Cre-mediated recombination between the *lox* sites. All stocks are available upon request.

### Cuticle preparations

Cuticle preparations were carried out as described in (Postika et al., 2018).

## Acknowledgements

We thank Farhod Hasanov for fly injections, Kate O’Connor-Giles for *pHD-DsRed* plasmid (Addgene plasmid # 51434), Bloomington Drosophila Stock Center for Drosophila lines.

## Author contributions

P.G., O.K. designed experiments. O.K., N.P. performed experiments. P.S., P.G., O.K. wrote the main manuscript text. O.K. prepared figures. All authors reviewed the manuscript.

## Competing interests

The authors declare no competing interests.

## Funding

This work (all functional and morphological analysis) was supported by the Russian Science Foundation, project no. 19-14-00103 (to O.K.). Part of the genome editing procedure was supported by grant 075-15-2019-1661 from the Ministry of Science and Higher Education of the Russian Federation. PS would like to acknowledge support from NIH R35 GM126975. Funding for open access charge: Russian Science Foundation.

## Notes

### Competing Interest Statement

The authors have declared no competing interest.

